# Wide genome transcription and metabolome profiles reprogrammed by sucrose under xenobiotic constraints

**DOI:** 10.1101/2022.05.31.493032

**Authors:** Richard Berthomé, Anne-Sophie Dumas, Emilie Jarde, Nataliia Ryzhenko, Evangelos Barbas, Abdelhak El Amrani

**Author notes:** Corresponding author: Abdelhak El Amrani.

## Abstract

Plants have developed strategies to adapt quickly to environmental changes. However, the regulation of these adaptive responses and coordination of signals network remains poorly understood for many environmental constraints. Indeed, signalling molecules play a central role in environmental stimuli and may coordinate plants development under environmental cues. Sucrose is the major transport carbohydrate in higher plants, in addition to its metabolic role; recent investigations suggested that sucrose impact the plasticity of plant development by controlling gene expression. Sucrose was also suggested as a ROS scavenger and as a signalling molecule. In this paper we showed that sucrose mitigated phenanthrene induced stress. Since, exogenous sucrose allowed growth and chlorophyll accumulation in the presence of high phenanthrene concentrations, whereas Arabidopsis development was blocked and seedlings were unable to accumulate chlorophyll in the presence of phenanthrene solely. To decipher the earlier molecular changes involved in sucrose tolerance to phenanthrene, wide transcriptional genes analysis and targeted metabolic profiling were carried out. We showed that sucrose driving plant response was associated with a deep reconfiguration of both genes expression and metabolites accumulation during the first hours of treatment, suggesting that sucrose, allowed plants to sustain the primary metabolism and to overcome the phenanthrene induced injuries.

## 1. Introduction

Contrary to animals, due to their sessile status higher plants had developed during their evolution, a larger plasticity response to environmental stresses. Consequently, their survival depends of their ability to escape and mitigate biotic and abiotic stress. Extensive investigations unraveled complex network signaling pathways which are involved in plant response to challenging environment, they showed the involvement of many common responses such as the induction of a set of stress related genes, ROS scavenging, and increase the synthesis of several metabolites. Soluble sugars accumulate under different abiotic stresses. Disaccharides such as sucrose, raffinose family polysaccharides (RFO) and fructans are water-soluble carbohydrates known to be involved in plant stress response (Rosa *et al*., 2009; Valluru and Ende, 2008). Soluble sugars have a central position in the cellular redox balance system: they are at the junction of photosynthesis, mitochondrial respiration and the beta-oxidation of fatty acids, pathways that lead to ROS production (Couée *et al*., 2006). An emerging concept suggested that sugars act as antioxidants (Ende and Valluru, 2009; Keunen *et al*., 2013) They are potentially involved in the modulation of oxidative stress and might interact with ROS signaling pathway (Bolouri-Moghaddam *et al*., 2010). In vitro studies have shown that the in vivo antioxidant capacity of soluble sugars have been underestimated: among other sugars, sucrose have been suggested to function as an antioxidant (Ende and Valluru, 2009; Bolouri-Moghaddam *et al*., 2010; Keunen *et al*., 2013). Several studies showed that sugars modulate a broad range of genes involved in central cellular processes: carbohydrate and nitrogen metabolism, signal transduction, metabolite transport and stress responses (Price *et al*., 2004; Thum *et al*., 2004).

Sucrose is considered as the major form of carbohydrate transport and the main product of photosynthesis in many plant species. It play a pivotal role in plants responses to abiotic stresses, and acts not only as a metabolite, but has also signaling properties (Koch, 2004; Wind *et al*., 2010; Rosa *et al*., 2009).

Sucrose has been shown to increase tolerance to a variety of abiotic stress such anoxia (Loreti *et al*., 2005), herbicide induced stress. Indeed, plants grown on sucrose showed increased tolerance to the herbicide (Sulmon *et al*., 2006; Ramel *et al*., 2007; Ramel *et al*., 2009). Hence, Sucrose enhances protection against reactive oxygen species (ROS) through the activation of specific ROS scavenging systems. This response is specific to sucrose as glucose and fructose exhibited lesser protection level.

Abiotic stresses influence plant development in natural ecosystems, among these challenging constraints, plants have to survive to the presence of a myriad of organic molecules which are mainly allelopatic signals, most of them are aromatic compounds. However, during the last decades anthropic activity have generated many aromatic end products such as polycyclic aromatic hydrocarbons (PAHs) which accumulated dramatically in natural ecosystems. Hence, environmental pollution constitutes a new world-wide ecological problem and may affect plant development.

Among the main pollutants, Polycyclic Aromatic Hydrocarbons (PAHs) are ubiquitous organic pollutant found in all the compartments of the environment. They are mainly considered as byproducts of anthropogenic activities with the incomplete combustion of petroleum products for example. This pollutant represents a great hazard for human health, creating the necessity to remediate contaminated environment.

Traditional clean-up techniques like incineration or physical-chemical processes represent a high cost and could also impact the environment. Indeed, development of green technologies using natural capacity of leaving organisms to store and degrade pollutants led to the creation of biological remediation techniques such as bioremediation and phytoremediation. Phytoremediation is the utilization of plants and their associated micro-organisms to decontaminate a polluted environment. This system has been proven efficient to eradicate PAHs contamination with several plant species. A model of plant detoxification system, inspired by the detoxification system of the mammalian liver, was proposed by Sandermann (1992) and named the “green liver model”. In other hand, Edwards *et al. (*2005) defined the xenome as the whole expressed genome involved in the detection, signaling, transformation and transport of xenobiotics. While little is known about the sensing and transport mechanisms, the “green liver” is a molecular model of the transformation and transport. Several multigenic enzymes families are involved in these processes: ATP-binding cassette transporter (ABC transporter), cytochrome P450 (CYP), Glutathione-S-transferase (GST) and UDP-glucuronosyltransferase (UGT).

It is of high interest to get a better understanding on how plants cope with the presence of these molecules in their environment. In-vitro studies, using the model plant *Arabidopsis thaliana*, revealed that phenanthrene, used as a model molecule of PAHs, cause several damages in plants. After a long-term (14 to 30 days) phenanthrene treatment, Arabidopsis presents a decrease in shoots development, roots length, chlorophyll content in a dose dependent way (Alkio *et al*., 2005; Liu *et al*., 2009).

Phenanthrene induced, like several abiotic stresses, an accumulation of reactive oxygen species (ROS). It also increased the activity of several anti-oxidant enzymes like the superoxide dismutase (SOD), the peroxide dismutase (POD) and the ascorbate peroxidase (APX) (Liu *et al*., 2009). These biochemical data were concordant with a transcriptomic analysis carried out in phenanthrene containing medium (Weisman *et al*., 2010) which revealed that phenanthrene induced significant change in genes expression involved ROS scavenging pathways.

The aim of this work was to examine if sucrose plays a protective effect when Arabidopsis was challenged with phenanthrene stressed condition, and to elucidate the early molecular events involved in the presence of exogenous sucrose. Transcriptional profiling and targeted metabolomic approaches were used to decipher the main signaling and metabolic pathways involved in sucrose induced tolerance to phenanthrene injury.

## 2. Material and methods

### 2.1. Plant material and growth conditions

Seeds of *Arabidopsis thaliana,* ecotype Columbia-0 (Col-0) were used in all experiments. Seeds were surface sterilized and sown on Petri dishes containing half-strength Murashige and Skoog (1962) medium, supplemented with sugar (sucrose, glucose or mannitol), phenanthrene and dimethylsulfoxide (DMSO).

As phenanthrene solubility in water is low, a solution of phenanthrene in DMSO was prepared at the concentration of 700mM and used to adapt the final concentration in the culture medium. DMSO was added in the same amount in all the conditions of each experiment. So that phenanthrene dose was the only parameter changing.

Petri dishes were placed at 4°C during 48h in order to break dormancy and to homogenize germination. Plants were grown at 22°C under a 16h-light period at approximately 5000lux.

### 2.2. Measurement of plant growth and development

For root measurement and determination of the fresh weight, plants were grown on MS/2 solid medium supplemented with 88mM of sugar (mannitol, sucrose or glucose), phenanthrene and methanol.

Fresh weight was determined after 22 days of growth for plants grown on MS/ supplemented with mannitol or sucrose at 88mM and with 0, 50, 100, 200 and 500µM of phenanthrene or with DMSO alone. Plantlets were grown on circle dishes horizontally for a better development and aerial parts were separated and harvested. For the two biological replicates, for each condition, 6 samples with 3 plantlets were harvested.

For a better observation of the primary root, plants were cultivated on Petri dishes vertically. Roots were measured after 14 days of growth on digital photographs using ImageJ v 1.45s software (Abramoff *et al*., 2004). For each condition, 30 independent plants were measured.

Results represent the mean with the standard error (SE). Statistical analyses were carried on using the Wilcoxon.test by R software (R development core team, 2010).

### 2.3. Fluorescence microscopy

Microscopic observations were performed at the Imagif platform, in the Cellular Biology Pole (CNRS, Gif-sur-Yvette, France).

Arabidopsis plantlets used in this experiment were grown on half-strength MS medium for 15 days, then transferred on half-strength MS medium supplemented with sucrose at 88mM containing 200µM of phenanthrene (prepared from a 700mM solution of phenanthrene in DMSO) or the same volume of DMSO. In order to facilitate the transfer, plants were grown vertically. A sterile and transparent plastic film was added to protect the shoot, so only roots are in contact with the contaminated medium.

After 5 days of phenanthrene treatment, leaves and roots were observed with a Zeiss LSM510 META microscope under UV-light (excitation 364nm and acquisition with 32 channels between 362nm and 704nm). Data were acquired using Zen2008 software developed by Zeiss (Germany).

### 2.4. Phenanthrene quantification

Determination of phenanthrene concentration in plants was performed in collaboration with Geosciences-UMR 6118 (Rennes, France).

Plants used for phenanthrene quantification were grown for 15 days on MS/2 then transferred in liquid medium supplemented with 0mM or 88mM of sucrose and containing 200µM of phenanthrene.

After 24h of incubation, plants were harvested and rinsed with water, absolute ethanol and then water. 3 samples of 20 plantlets pooled were taken. Plants samples were dried then pounded and weighted.

Phenanthrene was extracted from plants tissues by an accelerated solvent extractor (ASE 200, Dionex) with dichloromethane at 100°C and under a pressure of 100bar. Extracts were dried under a gentle flux of nitrogen, and finally weight for quantification using internal standard. One microliter of the extract was injected onto a Shimadzu QP2010+MS gas chromatograph/mass spectrometer (Shimadzu, Tokyo, Japan). The injector used was in splitless mode and maintained at a temperature of 310C. The chromatographic separation was performed on a fused silica SLB-5 ms capillary column (from Supelco, length = 60 m, diameter = 0.25 mm, film thickness = 0.25μm) under the following temperature program: 70°C (held for 1 min) to 130 at 15°C/min, then 130 to 300°C (held for 15 min) at 3°C/min. The helium flow was maintained at 1mL/min. The chromatograph was coupled to the mass spectrometer by a transfer line heated to 250°C. The analyses were performed in SIM mode (selective ion monitoring). Quantification was based on the internal standard phenanthrene-d10, which was added to the sample post-extraction and prior to the analysis by GC-MS.

### 2.5. Targeted analysis of metabolites

Analysis of metabolites levels were performed by the CORSAIRE platform (Biogenouest, INRA UMR 1359, Le Rheu, France).

Arabidopsis plants used for this analysis were grown on half-strength MS medium for 15 days, and then transferred on liquid half-strength MS medium supplemented with 0 or 88mM of sucrose and containing 200µM of phenanthrene (prepared from a 700mM solution of phenanthrene in DMSO) or the same volume of DMSO.

After 24 hours of incubation, plants were harvested, frost in liquid nitrogen, lyophilized and ground. A total of 10mg of dry plant material was extracted in 500µL of extraction solvent and 250µL of chloroform. The extraction solvent is composed of 5% of methanol and 95% of beta amino benzoic acid (10mM) – adonitol (20mM) concentrated solution. Samples were shaken 10 min at room temperature then 500µL of ultra pure water were added. All samples were vortexed for 30s then centrifugated for 5 min at 12000g at 15°C. The entire liquid phase is transferred to a new tube. For amino acids analysis, 50µL of the extract was dried under vacuum and 50µL of water were added. Samples are derivated using the AccQTag method (Waters) and analyzed by Ultra Performance Liquid Chromatography (UPLC, Waters). For sugars, organic acids, alcohol and ammonium quantification, 50µL of the extract supplemented with 50µL of internal standard are dried under vacuum. 50µL of methoxamine in pyridine (concentration 20mg/mL) were added to the dried samples which are incubated 90min at 30°C. 50µL of MSTFA (N-Methyl-N-(trimethylsilyl)trifluoroacetamide) are added to each sample which are incubated for 30min at 37°C and than analyzed by GC-MS.

### 2.6. Transcriptome studies

Transcriptome analysis was carried out at the Research Unit in Plant Genomics (INRA, Evry, France), using the CATMA version 5 microarrayscontaining 31776 specific gene tags corresponding to 22,089 genes from Arabidopsis (Crowe *et al*., 2003; Hilson *et al*., 2004).

*Arabidopsis thaliana* ecotype Columbia (Col 0) was grown *in-vitro* for 15 days on solid half-strength Murashige and Skoog (MS) medium. Plantlets at stage 1.04 (Boyes *et al*., 2001) were then transferred on liquid half-strength MS medium containing 0.2mM of phenanthrene (prepared from a 700mM solution of phenanthrene in DMSO) or the same amount of phenanthrene and 3% sucrose. Total RNA extractions of two independent replicates were performed using the Qiagen RNAeasy plant minikit according to the manufacturer’s instructions. Each biological replicates content phenanthrene-treated and phenanthrene plus sucrose treated plants. Each sample corresponding to 30 plants pooled were harvested after 30 min, 2h and 8h of incubation. For all comparisons performed, the experiment was done using the dye-switch technique. Lurin et al. (2004) described the protocol used for the labelling of antisense amplified mRNA with Cy3-dUTP and Cy5-dUTP (Perkins-Elmer-NEN Life science products), the hybridization to the slides and the scanning.

### 2.7. Statistical Analysis of microarray data

The Bioinformatic and Predictive Genomics group at the Research Unit in Plants Genomics (Evry, France), with whom the experiments were designed, developed specific statistics to analyse CATMA hybridations. For each array, the raw data include the logarithm of median feature pixel intensity (in log base 2) at wavelengths of 635 nm (red) and 532 nm (green). No background was subtracted. The normalization method used was described by Lurin *et al*. (2004). Differentially expressed genes were determined by performing a paired t-test on the log ratios averaged on the dye switch. A trimmed variance was calculated from spots that did not display extreme variance. The raw P values were adjusted by the Bonferroni method, which controls the family-wise error rate (with a type I error equal to 5%). We also adjusted the raw P values to control a false discovery rate using Benjamini-Yetkutieli at a level of 1%. Nonetheless, in the CATMA analysis pipeline, family-wise error rate proved to be the best solution to balance the estimated number of false positives and false negatives (Ge *et al*., 2003). As described by Gagnot et al. (2008), when the Bonferroni P value was lower than 0.05, the gene was declared differentially expressed.

### 2.8. Biological pathways enrichment

Biological pathways significantly over-represented in the list of differentially expressed genes were identified with the classification superviewer tool of the university of Toronto website (http://bar.utoronto.ca/ntools/cgi-bin/ntools_classification_superviewer.cgi) using MAPMAN classification (Provart T., 2003) as a source.

### 2.9. Microarrays data validation by quantitative Real-Time PCR (qRT-PCR)

The qRT-PCR validation was carried on 15 genes being found differentially expressed in the microarrays experiments.

Primers were designed using the online software Primers3 ((Rozen and Skaletsky, 2000), http://frodo.wi.mit.edu/, optimal temperature 60°C, Supplemental Table S2).

The primer pairs were first tested on a dilution series of genomic DNA (5, 0.5, 0.05, and 0.005ng) to generate a standard curve and assess their PCR efficiency, which ranged between 90% and 110%. RT was performed on 1µg of total RNA with oligo(dT) primer (18-mer) and the SuperScript II RNase H^-^ reverse transcriptase according to the manufacturer’s instructions (Invitrogen). In every run, at least three replicates PCRs were done for each cDNAs. For each gene investigated using qPCR, a dilution series covering 3 orders of magnitude (1, 1/10 and 1/100) of the cDNA stock solution was prepared. Three replicates of each of the three standards were used in qPCR experiment together with three no-template controls. qPCR was performed in 5 L, with 0.5 L of RT reaction (1/100 dilution), 900 nM final concentration of each primer pair, and 2,5µL of MESA GREEN qPCR MasterMix Plus for SYBR® Assay (Eurogentec)). Corresponding minus-RT controls were performed with each primer pair. All reactions were performed with the CFX384 Touch™ Real-Time PCR Detection System (Bio-Rad) as follows: 95°C for 5 min; 40×95°C for 10 s and 60°C for 30 s; and a dissociation step to discriminate primer dimers from the PCR product. Using CFX Manager™ Software version 3.0 provided by the manufacturer, the optimal cycle threshold (Ct) was determined from the dilution series, with the raw expression data derived. Six housekeeping genes were assessed in this experiment, and the two best control genes, consistently expressed, were selected to calculate the average normalization factor: *AT4G13615* and *AT5G21090* for each sample pair. Normalized (Norm) ΔCt for each differentially expressed gene was calculated as following: Norm ΔCt = - (rawΔCt - Norm factor). Microarray data from this article were deposited at Gene Expression Omnibus (http://www.ncbi.nlm.nih.gov/geo/), accession no. GSE48181) and at CATdb (http://urgv.evry.inra.fr/CATdb/; Project: AU10-04_phytoremediation) according to the “Minimum Information About a Microarray Experiment” standards.

## 3. Results and discussion

### 3.1. Sucrose induced tolerance to phenanthrene injury

Sucrose was shown to protect the plant against abiotic stresses such anoxia and herbice injury (Loreti *et al*., 2005; Sulmon *et al*., 2004). However no data were available about its effect on stresses induced by organic pollutants such as PAHs. In this work, we studied the incidence of sucrose implementation on plant response to phenanthrene exposition.

Arabidopsis plants were grown on media containing 3% w/v of sucrose (=88mM), then compared to control plants grown in the same culture medium supplemented with mannitol, used as previously described as an osmoticum (Borsani *et al*., 2001; Sulmon *et al*., 2004). In these conditions, plants were submitted to a phenanthrene treatment at different concentrations ranging from 0µM to 400µM. Shoots fresh weight was measured as marker of development to assess sucrose effects. Control plants, grown on mannitol supplemented medium, showed a significant reduction of growth development even at low phenanthrene concentration (50 µm) (Fig.1 and Fig.2), and reduction of the number and leaves surface. 70% of plants failed to accumulate chlorophyll (Fig.1). In parallel, when phenanthrene increased up to 100 µm, 100% of plants showed high toxicity symptoms and are unable to initiate first leaves and seems to be unable to initiate chlorophyll synthesis. On the contrary, plants growing on sucrose supplemented media showed a significantly highest fresh weight compared to plants grown on mannitol-supplemented media whatever the phenanthrene concentration applied, and even at high phenanthrene concentration (400 µm) plants maintained growth and developed leaves and chlorophyll biosynthesis.

**Figure 1:**
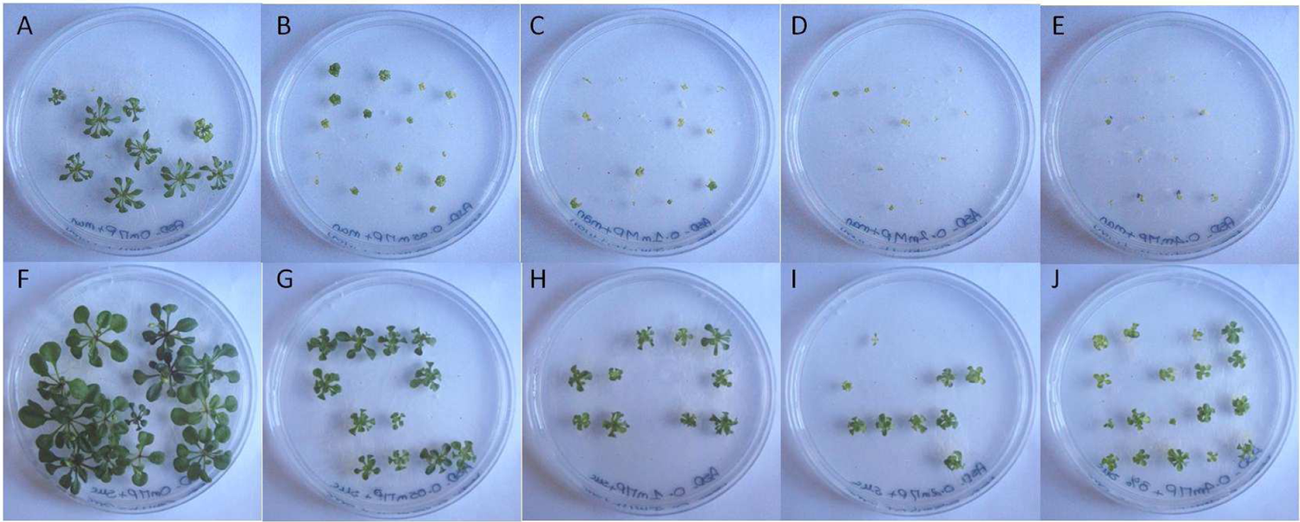
Sucrose mitigated phenanthrene induced stress. 22 days old plantlets were grown on half MS medium. Phenanthrene was supplemented at 0µM (A, F), 50µM (B, G), 100µM (C, H), 200µM (D, I) and 400µM (E, J). Control plants (A to E) were grown on medium supplemented with mannitol as an osmoticum, plants from pictures F to J grown on sucrose supplemented media.

**Figure 2:**
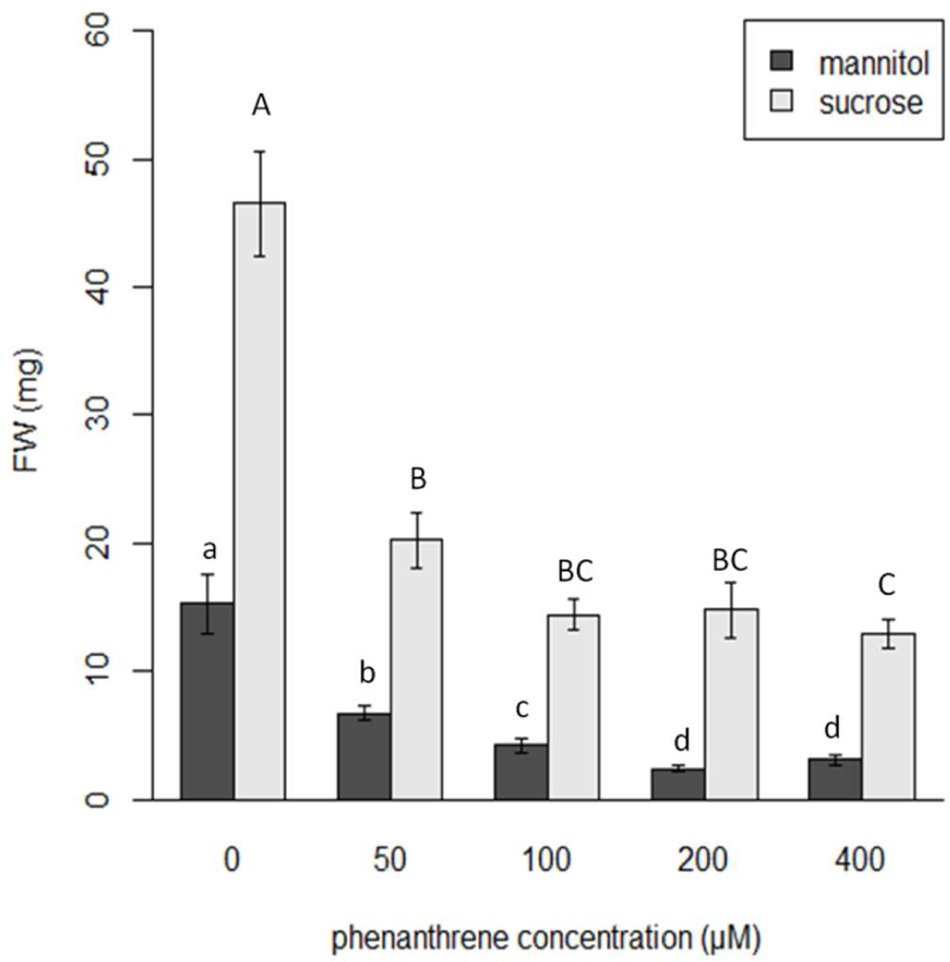
Fresh weight of 22-day-old plants following a phenanthrene treatment (0µM, 50µM, 100µM, 200µM and 400µM) in a growth medium supplemented with sucrose or mannitol (used as osmoticum). Different letters indicate statistical difference (p-value≤0.05) between doses of phenanthrene.

In order to assess the specific sucrose effect and to better understand the role of sucrose in plant development under a phenanthrene-induced stress, a comparison has been carried out with plants grown on media supplement with glucose. Both molecules were exogenously added to the growing medium at 88mM under high concentration of phenanthrene ranging from 200µM to 1000µM. In this experiment, we used primary root elongation as growth marker, since this later was highly sensitive to abiotic stresses (Fig. 3). Statistical analysis showed that sucrose induced a pronounced effect in high phenanthrene supplemented medium, even if glucose mitigated the phenanthrene inhibition of the primary roots, this protection was at a lesser extent. This observation corroborates with several already published data about the specific effect of sucrose. Solfanelli *et al*, (2006) carried out transcriptome analysis and showed that sucrose exhibited specific induction of the anthocyanin biosynthetic pathway, suggesting the involvement of specific sucrose signaling pathway. Emerging idea suggested that sucrose impacts also plant tolerance to environmental stress. Loreti et al (2010) showed exogenous sucrose greatly enhances anoxia tolerance of Arabidopsis seedlings but glucose did not substitute for sucrose in this process. Sucrose exhibited similar effect when Arabidopsis was challenged with atrazine. In these conditions sucrose induced the ROS scavenging system and allowed plant development and growth (Sulmon *et al*., 2006; Ramel *et al*., 2007).

**Figure 3:**
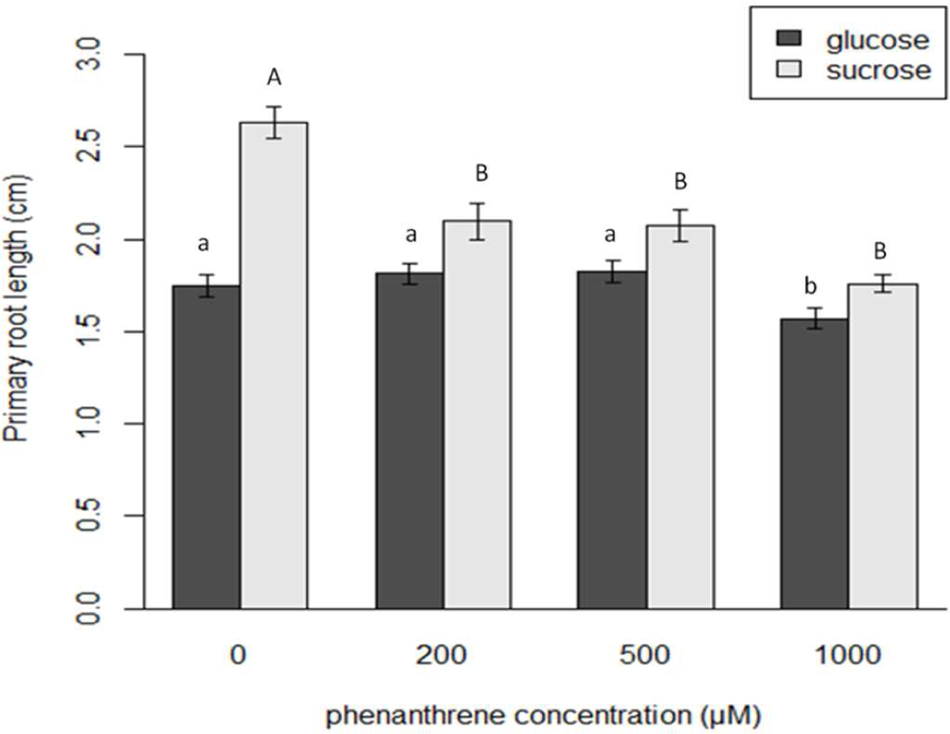
Primary root length of 14-day-old plantlets after a phenanthrene treatment (grown on 0µM, 200µM, 500µM and 1000µM respectively) in a growth-medium supplemented respectively with 88mM of sucrose or glucose. Small letters show differences between different phenanthrene doses for plants treated with glucose while capital letters show differences between different phenanthrene doses for plants treated with sucrose. (p-value≤0.05)

Indeed, Barker et al. (2000) proposed that sucrose transporter SUT2 act as a sucrose sensor, but this assumption, has not been demonstrated yet. Since the elucidation sucrose-specific signaling pathway is difficult to study as sucrose is readily hydrolyzed into hexoses. However, functional genomics and omics approaches should give new insight to understand how this important molecule controls plant development and tolerance to environmental constraints.

### 3.2. Cellular and histological localization of phenanthrene under protective sucrose condition

It is of high interest to understand how sucrose modulates the phenanthrene accumulation in plant cells and tissues. In this paper, we used fluorescence properties of free phenanthrene under UV light (Dabestani and Ivanov, 1999) to study its *in*-*planta* localisation. Indeed, after an excitation at 564nm, a specific signal was emitted between 450-500nm, corresponding to phenanthrene fluorescence. At least, these observations enable us to localize native phenanthrene within leaving plant cells.

In sucrose supplemented medium, confocal microscopy analysis showed accumulation of free phenanthrene in the trichome (Fig.4A) and in the abaxial leaf surface (Fig.4E). In addition, sucrose seems to allow the accumulation in vascular tissues (Fig. 4I). This last result was different from those published in Dumas et al. (2016). Showing that phenanthrene accumulated only in trichome and abaxial cells of Arabidopsis leaves, when plants were grown without exogenous sucrose.

**Figure 4:**
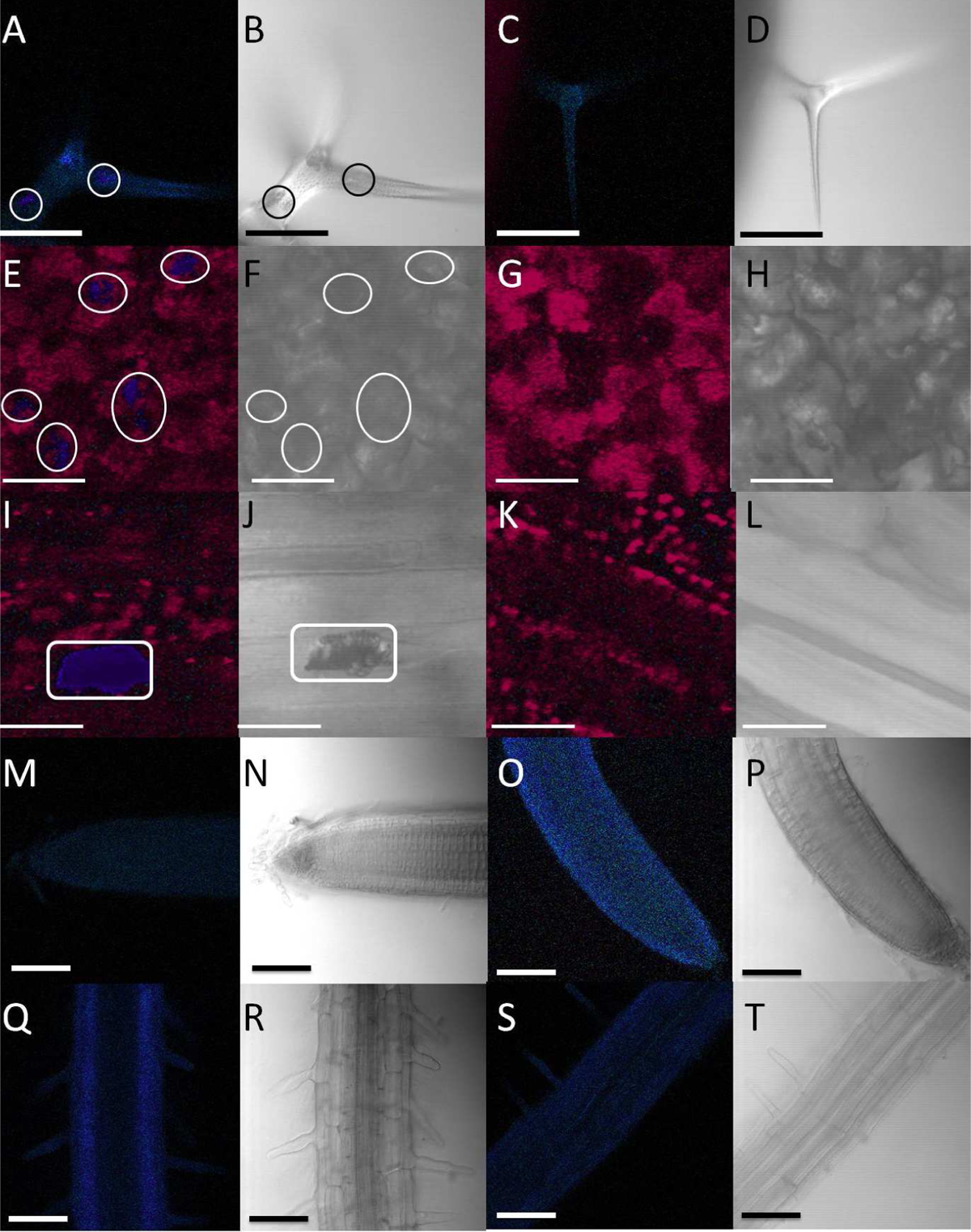
Cellular and histological localization of free phenanthrene. Plants were treated for 6 days with 0.2mM of phenanthrene + sucrose (A, B, E, F, I, J, M, N, Q and R) or DMSO + sucrose as control (C, D, G, H, K, L, O, P, S and T). Pictures A to D are an observation of the trichomes, E to H of the underside of the leaves, I to L of the vein from the underside of the leaf, M to P of the apex of the primary root and Q to T of the root hairs of the primary root. 2 consecutives photos (A-B, C-D …) represent the same area. A, C, E, G, I, K, M, O, Q and S are observations made under UV light and B, D, F, H, J, L, N, P, R and T with transmission. Circles show the localization of phenanthrene. All bar scales represent 100µm except for pictures E to L where it represents 50µm.

These results corroborate data published by Alkio et al. (2005) who showed that plant treated with phenanthrene during 30 days on a sucrose containing medium harbor local necrosis similar to hypersensitive responses. These authors also found that the frequency and size of these lesions were phenanthrene concentration-dependent.

These results suggested that sucrose triggered discrete changes in the cellular and tissue transport and/or accumulation of the free phenanthrene, as this later was also localized in vascular tissues.

In order to quantify phenanthrene accumulation in plant tissues, 16 days old plantlets were transferred to liquid medium containing phenanthrene alone (control) or phenanthrene plus sucrose. After 24 h of incubation, phenanthrene quantification was carried out and surprisingly showed that plants grown on sucrose supplemented medium accumulated 50% less phenanthrene than plants grown on a sugar free medium (Fig. 5). This suggested that sucrose induced either new gene-network involved in phenanthrene metabolization, transport or likely a complete degradation, which were silenced when phenanthrene was present alone. All these data taken together suggested that sucrose reconfigure cellular and molecular changes involved in storage and metabolization strategy to induce tolerance to phenanthrene injury.

**Figure 5:**
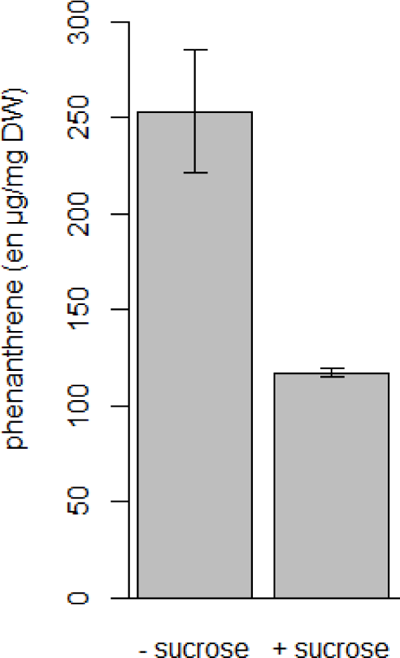
Phenanthrene content of 16-day-old plantlets incubated for 24h with 200µM of phenanthrene, supplemented with or without sucrose in the growth medium.

### 3.3. Sucrose reconfigured transcriptomic profile under phenanthrene induced stress

#### 3.3.1. General effects of sucrose under phenanthrene induced stress

Transcriptional analysis was carried out at the developmental stage 2.04 (Boyes *et al*., 2001). Plantlets were grown on solid medium for 15 days, then were transferred respectively into liquid medium supplemented with 200µM of phenanthrene or with 200µM of phenanthrene in protective condition (88mM of sucrose), during the incubation time liquid medium was agitated in order to avoid any oxygen deprivation.

Transcriptional analysis was performed using CATMA microarrays (version 5) and two independent biological replicates were made. The experimental design was designed to study the sucrose effect on transcriptional response of plants incubated with phenanthrene-during 0.5h, 2h and 8h. We will refer this experiment as the “sucrose experiment”, in contrast to the microarray experiment previously performed to study the plant response to phenanthrene alone (Dumas et al. (2016) that will be referred as the “phenanthrene experiment”. Overall, 2088 genes were differentially expressed at least one time in one condition of the “sucrose experiment” (DEG). The numbers of DEG at each point of the kinetic is presented in figure 6. The comparison between number of DEG obtained at each time point, revealed a transcriptional regulation shift after 4h of phenanthrene incubation, with an important increase of DEG at 8h compared to 4h. These results suggest that plant responses to phenanthrene in the presence of exogenous sucrose might be divided into two steps: an early response up to 4H and a late response which started at 8h. 72% of the DEG at 0.5h and 66% of the DEG at 8h are specific to each kinetic point while only 28% are specific to the 2h incubation time. These results reinforce the idea that the 2h kinetic point seemed to be an intermediate between both stages.

**Figure 6:**
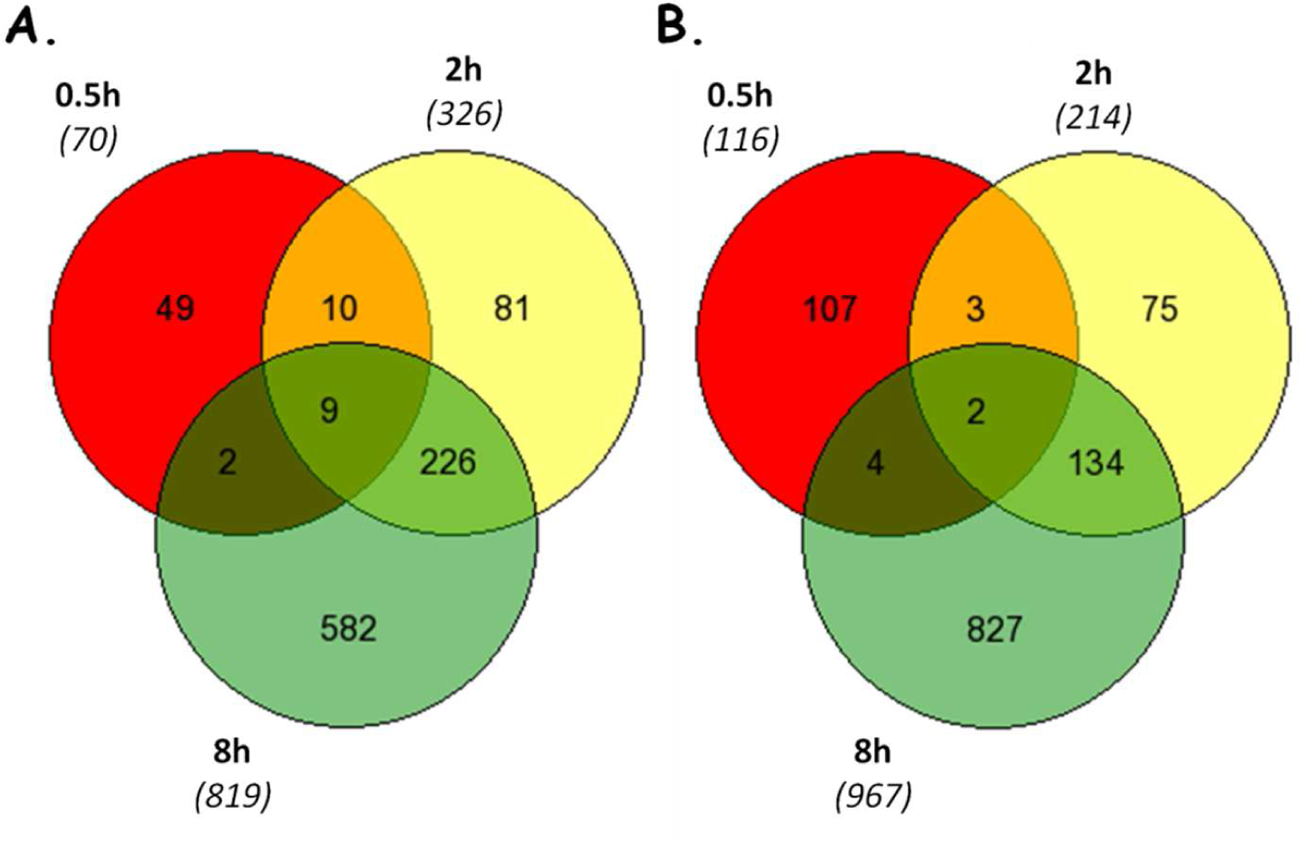
Number of down-(A.) and up-(B.) regulated genes at 0.5h, 2h and 4h Venn diagrams were generated using the online application Gene List Venn Diagram (http://simbioinf.com/mcbc/applications/genevenn/genevenn.htm, (Pirooznia et al., 2007))

The most affected pathways by sucrose under phenanthrene-induced stress were determined using the classification superviewer tool from the Bio-Array Resource for Plant Biology (Provart T., 2003) using the MapMan (Thimm *et al*., 2004) classification as the source.

Primary metabolisms (photosynthesis, carbohydrates metabolism, fermentation, glycolysis, N-metabolism…) were affected by sucrose, but also development, transport, RNA (transcription factors), secondary metabolism, signaling, protein metabolism (Fig. 7). As it could be expected, sucrose seems to modify primary metabolism relatively quickly. Most of these pathways are known to be regulated by sucrose (Coruzzi and Zhou, 2001; Price *et al*., 2004; Thum *et al*., 2004; Rosa *et al*., 2009).

**Figure 7:**
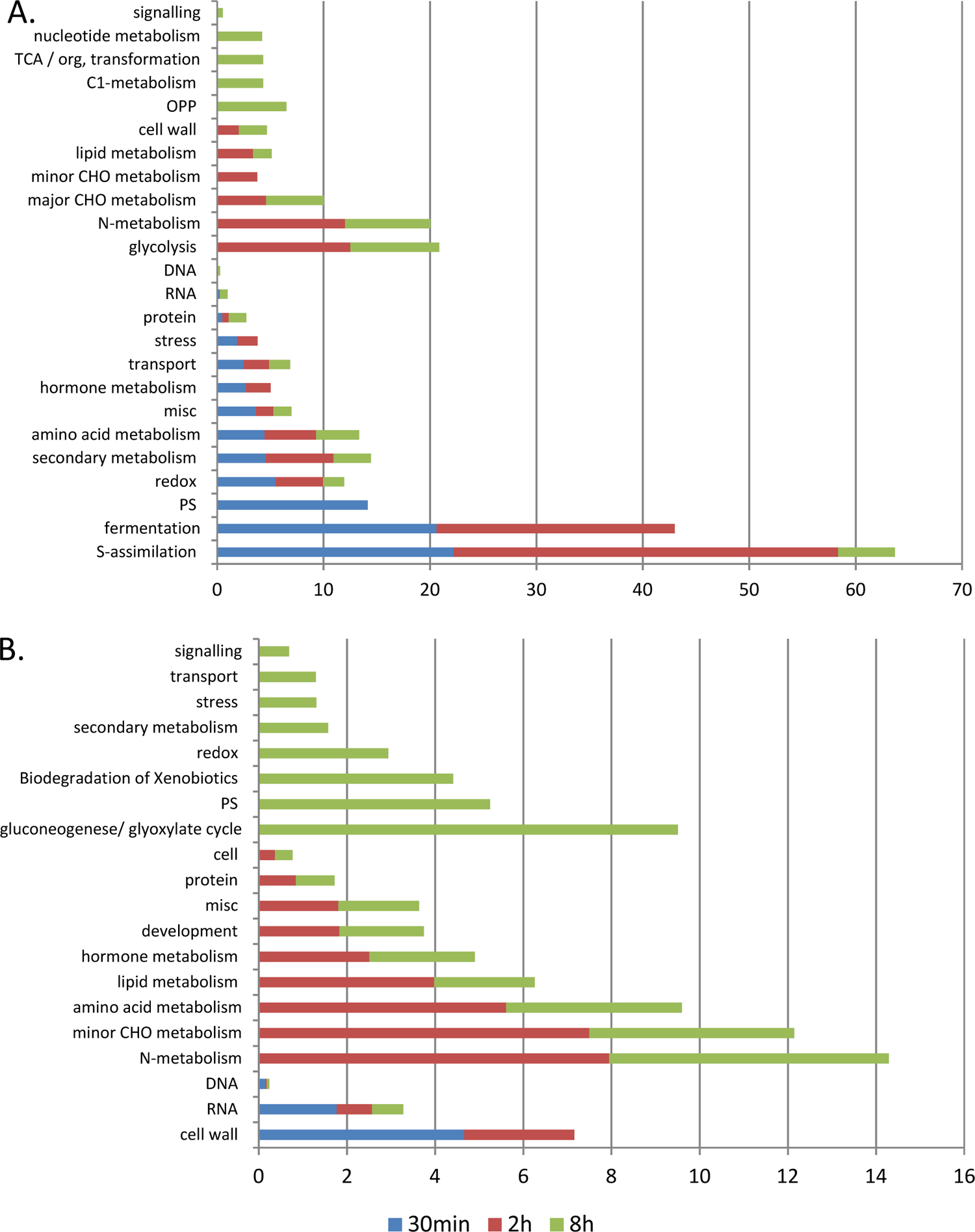
Biological pathways with significant over-representation of up-(A) and down-regulated (B) genes (P-values<0.05) after 0.5h, 2h and 8h of treatment with sucrose under phenanthrene-induced stress. Functional enrichment is shown for differentially expressed genes analyzed using the classification superviewer tool from the Bio-Array Resource for Plant Biology (Provart T., 2003) (http://www.bar.utoronto.ca/ntools/cgi-bin/ntools_classification_superviewer.cgi) using the MapMan classification as the source. PS, Photosynthesis. Misc, miscellaneous.

During the early stage, among the significant up regulated pathways by sucrose, we found a stimulation of photosynthesis, and strong regulation of genes involved in stress responses and red-ox scavenging system. The late stage showed up regulation of pathway mostly related to general sucrose responses, with a strong regulation of primary metabolism but also the stimulation of the redox system. Some pathways are overrepresented in both stages but the DEG belonging to those pathways are different according to the stage.

#### 3.3.2. Effect of sucrose on primary metabolism pathways under phenanthrene induced stress

Sucrose is the major form of transport of carbohydrates in plants, it is known that the primary metabolism is closely regulated by this molecule. Indeed, our transcriptional analysis showed that photosynthesis was affected rapidly by sucrose. After 30 minutes, photosynthesis showed quick and very significant induction. However, after 8 hours of most of these genes were down regulated. This repression of photosynthesis after 8 hours of treatment could be in accordance with an optimal utilization of the carbon source in the growth medium previously described (Roitsch, 1999; Rolland *et al*., 2002). However, the stimulation of photosynthesis in the early stages of the experimentation is surprising.

Meanwhile, glycolysis was up-regulated after 2 hours and 8 hours of treatment, while TCA was up-regulated only after 8 hours of treatment. Fermentation was up-regulated after 0.5h and 2 hours of treatment. This phenomenon was also observed in response to a 24 hours treatment of phenanthrene alone (see Dumas et al (2016). During our experimental design, attention have been carried out to avoid any oxygen deprivation. Hence, seedlings were incubated in liquid medium before being harvested for transcriptome analysis. However, in these conditions oxygen deprivation might occur. To exclude this possibility, we compared our gene lists to genes differentially regulated by anoxia, hypoxia, and O2 deprivation in the seedlings/shoots of Arabidopsis microarray datasets (55 genes) and showed that only 3 genes are found in common (see Dumas et al (2016).

Nitrogen metabolism is also affected by sucrose under phenanthrene induced stress after 2 and 8 hours of treatment. Glutamate and glutamine synthesis pathway is up-regulated at this stage. Sucrose induced a strong regulation of amino acids metabolism: amino acids synthesis is mostly enhanced while the degradation is mostly repressed. This strong regulation of these metabolic pathways is characteristic of the addition of exogenous carbon (Coruzzi and Zhou, 2001) and is also observed by Ramel et al. (2007) under atrazine-induced stress.

Data from the “phenanthrene experiment” (Dumas et al. 2016) revealed that phenanthrene strongly affected primary metabolism, as photosynthesis and respiration were dramatically down regulated. Meanwhile, soluble sugars and amino acids levels increased. This physiological unbalance revealed a carbohydrate starvation as cells are enable to face energetic demand. This unbalance was associated with chlorotic phenotype and inhibition of plant development. However, when sucrose is supplied, it seems to maintain metabolic pathways involved in the production of carbon skeletons (photosynthesis) and energy production (fermentation, glycolysis TCA cycle) (see fig: 7). It may be argued that sucrosecompensates the negative effect of phenanthrene as it is used as metabolic molecule to face energy demand since it might induce new xenome transcription changes in order to reduce phenanthrene toxicity or even metabolization. In order to understand how sucrose mitigates phenanthrene induced stress at the transcriptional level, all differentially expressed genes family belonging to the xenome were identified and compared with phenanthrene induced stress condition. This approach might bring new insight into sucrose.

#### 3.3.3. Phenanthrene induced oxidative stress genes was reconfigured by exogenous sucrose

Several published data showed that phenanthrene was known to induce oxidative stress (Alkio *et al*., 2005; Liu *et al*., 2009; Weisman *et al*., 2010). In parallel it might induce also several genes involved in detoxification at the wide genome level. It is of high interest to study how these specific genes family are regulated in the presence of sucrose.

Results obtain from pathways enrichment analysis (Fig. 7) revealed that up-regulated genes belonging to the stress response pathways were over-represented after 0.5h minutes and 2 hours of treatment while genes involved in these pathways were mostly repressed after 8 hours of treatment. Overall, most genes involved in these two stages of stress response are distinct. This result suggested that stress response genes are regulated differently according to the time course of incubation. Surprisingly, figure: 8 showed that genes of stress response are mostly down-regulated in the “sucrose experiment” while those genes were up-regulated in the “phenanthrene experiment”. Consequently, sucrose mitigated phenanthrene induced stress at the transcriptional level, as it reduces the transcriptional activity of significative expressed genes involved in stress response. The pathway corresponding to the DEG up-regulated of the redox system is significantly over-represented over the whole experiment (Fig. 7). However, the one corresponding to the DEG down-regulated is only over-represented after 8 hours of treatment. These results suggest that the regulation of the genes corresponding to the redox system are highly induced especially after 8 hours in phenanthrene treatment conditions.

Among the 31 genes differentially expressed belonging to this pathway, 13 are involved in the glutathione and ascorbate redox pathway. Phenanthrene has been described to increase the activity of anti-oxidant enzymes such as ascorbate peroxidases (APX) and catalases (CAT) (Liu *et al*., 2009). Sugars are often linked to the anti-oxidant system. *In-vitro* studies revealed that sucrose showed a strong antioxidant activity but these results have not been found *in-planta* yet (Ende and Valluru, 2009; Bolouri-Moghaddam *et al*., 2010). Moreover, Ramel et al. (2009) also showed that sucrose protection from atrazine injuries involved a strong regulation of the redox system. 7 of the 11 up-regulated genes were also up-regulated in the “phenanthrene experiment” (Fig. 9). Sucrose increased redox genes expression, which is concordant with the sensing role of sucrose on redox pathway. Data of Liu et al. (2009) were obtained by studying phenanthrene effects on plants grown on media supplemented with sucrose. These data highlighted that phenanthrene induced the activity of anti-oxidant enzymes. These results were corroborated by Weisman et al. (2010) transcriptomic analysis which revealed the stimulation of multiple genes involved in the redox system. At the contrary, data from the ‘phenanthrene experiment’ (Dumas et al (2016) did not show that oxidative stress was one of the main effect of phenanthrene on plants after a short term exposure. These results from multiple experiments suggested that the activation of the redox system under phenanthrene induced stress might be provoked by the exogenous sucrose. Finally sucrose seems to regulated, at the transcriptional level, genes involved in antioxidant response upon phenanthrene induced stress, revealing the specific effect of this molecule.

**Figure 8:**
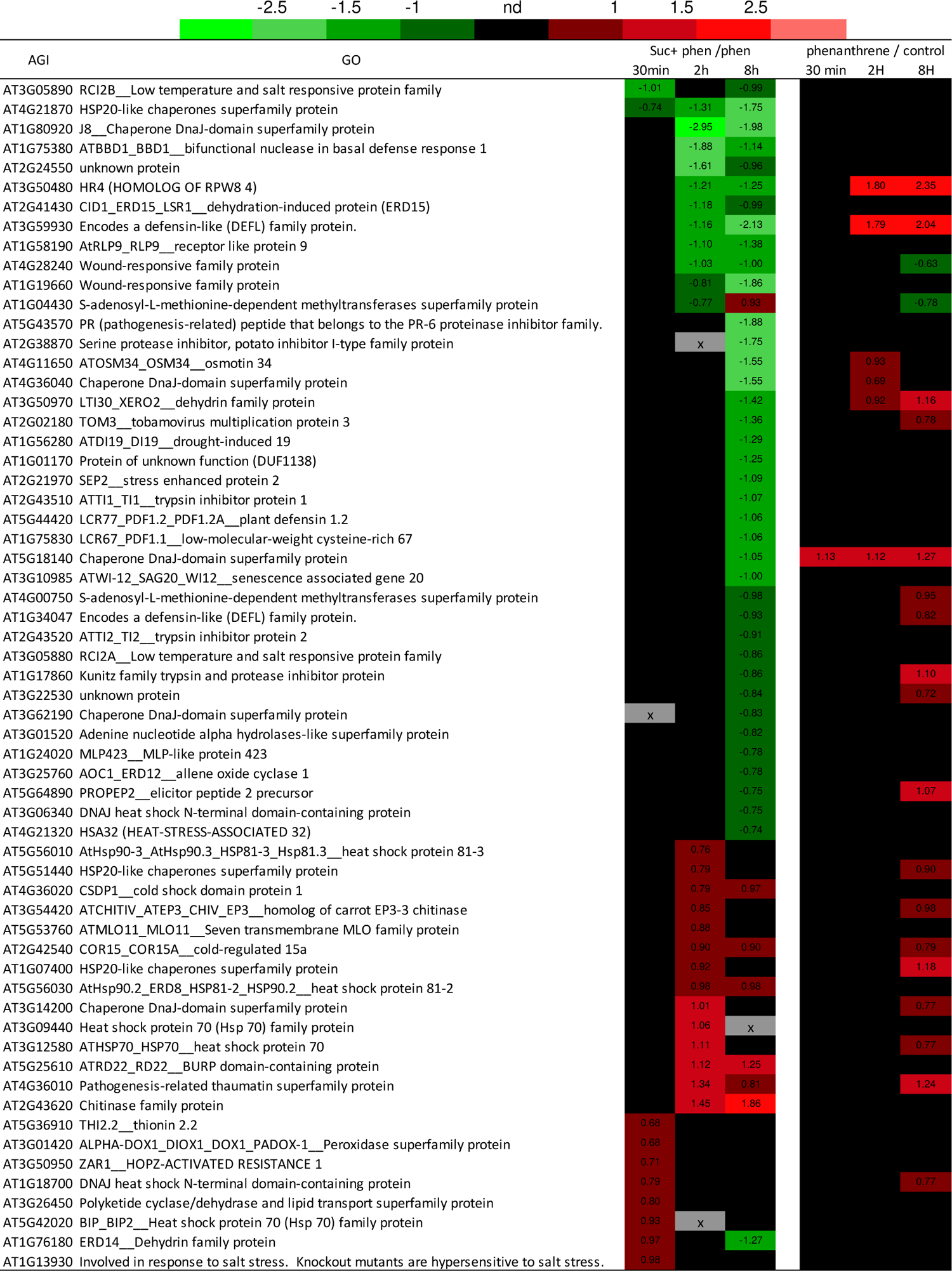
List of differentially expressed genes belonging to the stress response pathway in the “sucrose experiment”. In parallel, data extracted from the “phenanthrene experiment” were also shown (right) (Dumas et al (2013) submitted paper). A negative ratio indicates that the gene is decreased and a positive ratio indicates that the gene is increased in expression in the indicative point of the kinetic, respectively. Ratios in black boxes were not found to be statistically significant after Bonferroni correction (P >0.05)

**Figure 9:**
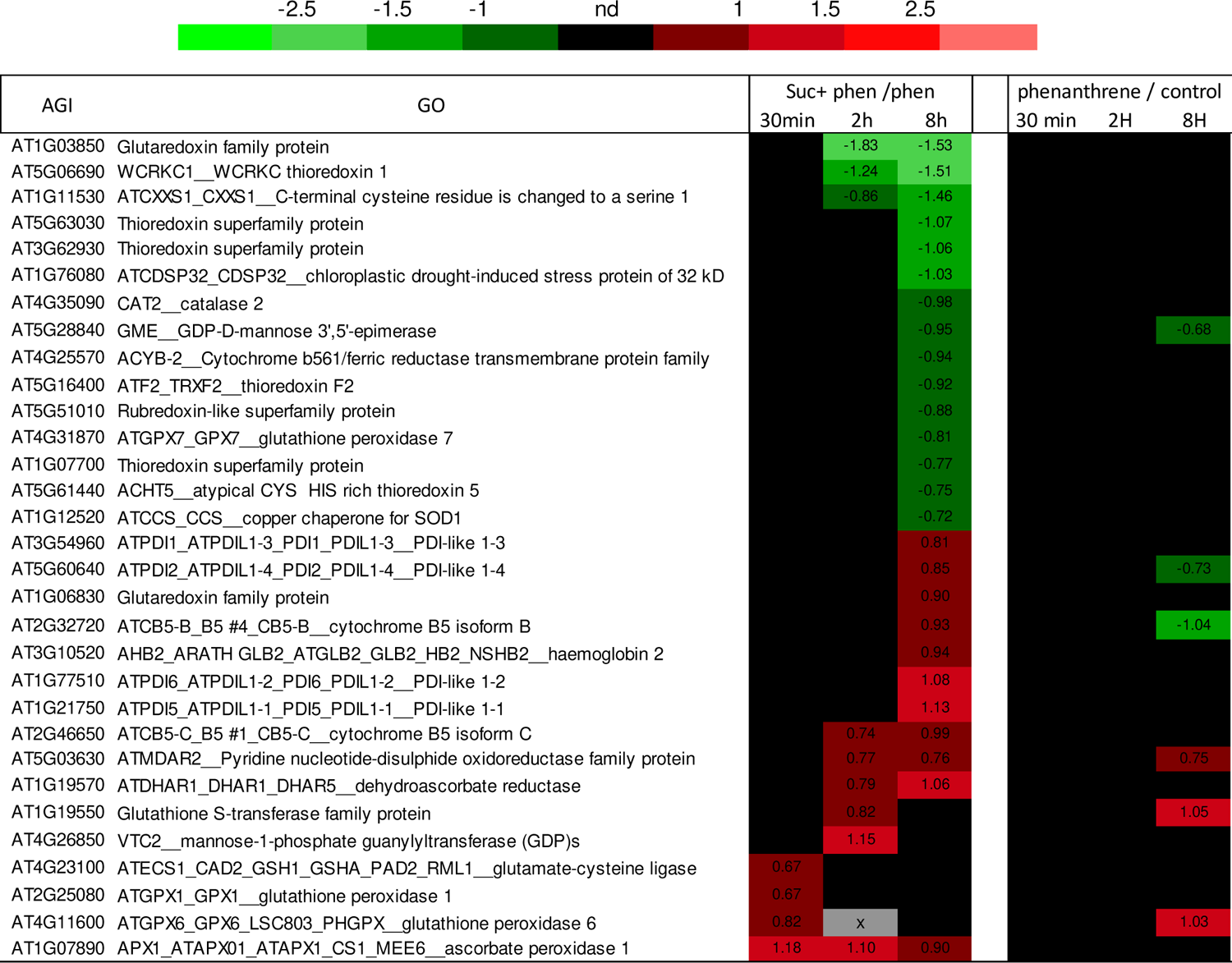
List of differentially expressed genes of the redox pathway in the “sucrose experiment”.In parallel, data extracted from the “phenanthrene experiment” were also shown (right) (Dumas et al (2013) submitted paper).A negative ratio indicates that the gene is decreased and a positive ratio indicates that the gene is increased in expression in the indicative point of the kinetic, respectively. Ratios in black boxes were not found to be statistically significant after Bonferroni correction (P >0.05)

#### 3.3.4. The xenome expression was reconfigured by exogenous sucrose

The transport and detoxification encoding genes of the xenome involve several multigenic enzymes families: ATP-binding cassette transporters (ABC transporters), glutathione-S-transferases (GST), UDP-glucuronosyltransferase (UGT) and cytochrome P450 (CYP) (Edwards *et al*., 2005). The complete list of xenome genes differentially expressed in this experiment is listed in figure 10. This list was confronted to results previously obtained in phenanthrene treated plants. Exogenous sucrose reduced the transcriptional activity of most (52 %) xenome genes which were induced in the presence of phenanthrene alone. Sucrose seems to reverse the effect of phenanthrene at the xenome level. Meanwhile, stimulates other xenome enzymes. Indeed, sucrose led to the activation of a new xenome expression.

**Figure 10:**
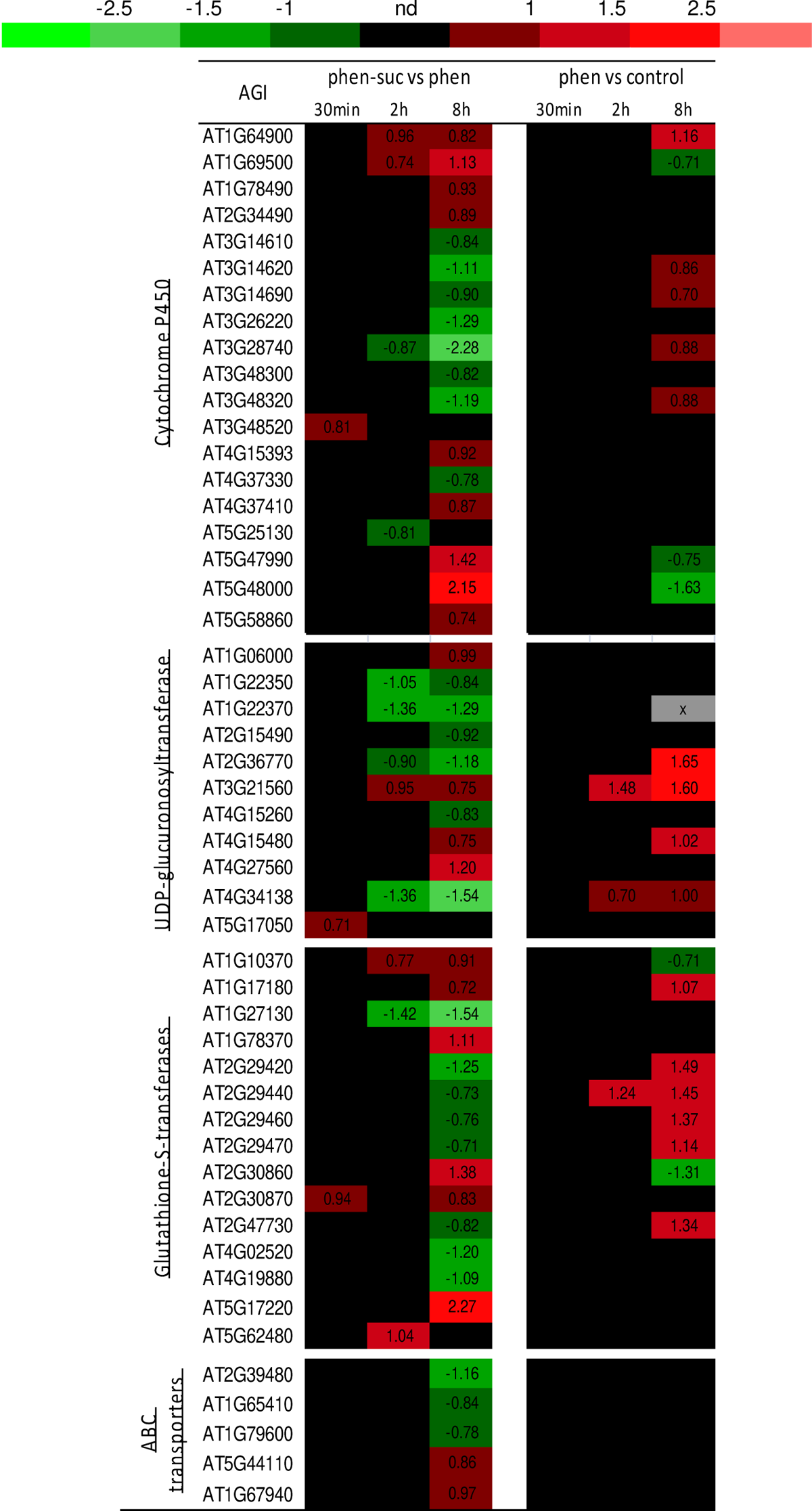
List of differentially expressed xenome in the “sucrose experiment”, in parallel, data extracted from the “phenanthrene experiment” were also shown (right) (Dumas et al (2016). A negative ratio indicates that the gene is decreased and a positive ratio indicates that the gene is increased in expression in the indicative point of the kinetic, respectively. Ratios in black boxes were not found to be statistically significant after Bonferroni correction (P >0.05)

#### 3.3.5. Confrontation with others experiments

Other microarrays experiments have been performed to study sucrose effects under atrazine stressed plants (Ramel *et al*., 2007) or under anoxia conditions (Loreti *et al*., 2005). While Ramel et al. used the CATMA technology as we did, Loreti utilized Affymetrix technologies. Among the 1967 DEG of our experiment, 1034 are also differentially expressed in at least one time in other experiments studying the sucrose effect. 4 genes are down regulated in all the experiments and none is up-regulated in all experiments.

Comparing the DEG of our experiment and the ones of the “control condition” experiment (sucrose vs. mannitol) (Ramel *et al*., 2007 and Loreti *et al*., 2005) should give us a list of genes specifically regulated in the presence of sucrose. Indeed, among the 738 genes that are common to both experiments, 655 were found to be DEG, identically repressed or enhanced in both conditions.

Using the classification superviewer tool from the Bio-Array Resource for Plant Biology (Provart T., 2003) with the MapMan (Thimm *et al*., 2004) classification as the source, pathways significantly over-represented in the 655 DEG were determined. Primary metabolism pathways like nitrogen metabolism, glycolysis, fermentation, pentose phosphate pathway (OPP) and photosynthesis are the most affected by sucrose. These genes represent 90% of the common genes revealing the effect of sucrose, mostly on in primary metabolism. This sucrose effect is not only due to the accumulation of exogenous sucrose in the plant but also to signaling effects.

In parallel, 288 genes are differentially expressed in our experiment and in a sucrose effects under abiotic stress experiment (Ramel *et al*., 2007 and Loreti *et al*., 2005). Pathway enrichment analysis revealed that these genes are mostly involved in carbohydrates metabolism, amino acids metabolism but also in redox system, secondary metabolism, development and stress response. These genes are belonging to pathways known to be regulated by sucrose under abiotic stress. These results revealed that sucrose effects under phenanthrene-induced stress are relatively close to sucrose effects under others abiotic stresses.

### 3.4. Targeted metabolome profiling under sucrose protective conditions

Metabolites analysis was carried out on plants at the stage 2.04 (Boyes *et al*., 2001). 15 days old plantlets were grown on solid basal medium, then transferred into liquid medium. The analysis was carried on as fallow: control plants grown in the basal medium (control), plants grown in medium with sucrose only (named sucrose), plants grown in the presence of phenanthrene only (named phenanthrene) and plants grown in the protective condition in the presence of sucrose and phenanthrene (named sucrose + phenanthrene).

Comparisons between these four conditions allowed us to determine metabolites changes under phenanthrene-induced stress in unprotected condition 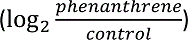 and how sucrose mitigated the effects of phenanthrene 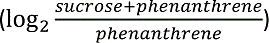. These results will be compared with changes induced by sucrose alone 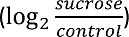. Among the 47 metabolites analyzed, 31 showed a significant change in their level at least one comparison (one ratio). Figure 11 summarized these results while complete data can be found in supplemental table S1.

**Figure 11:**
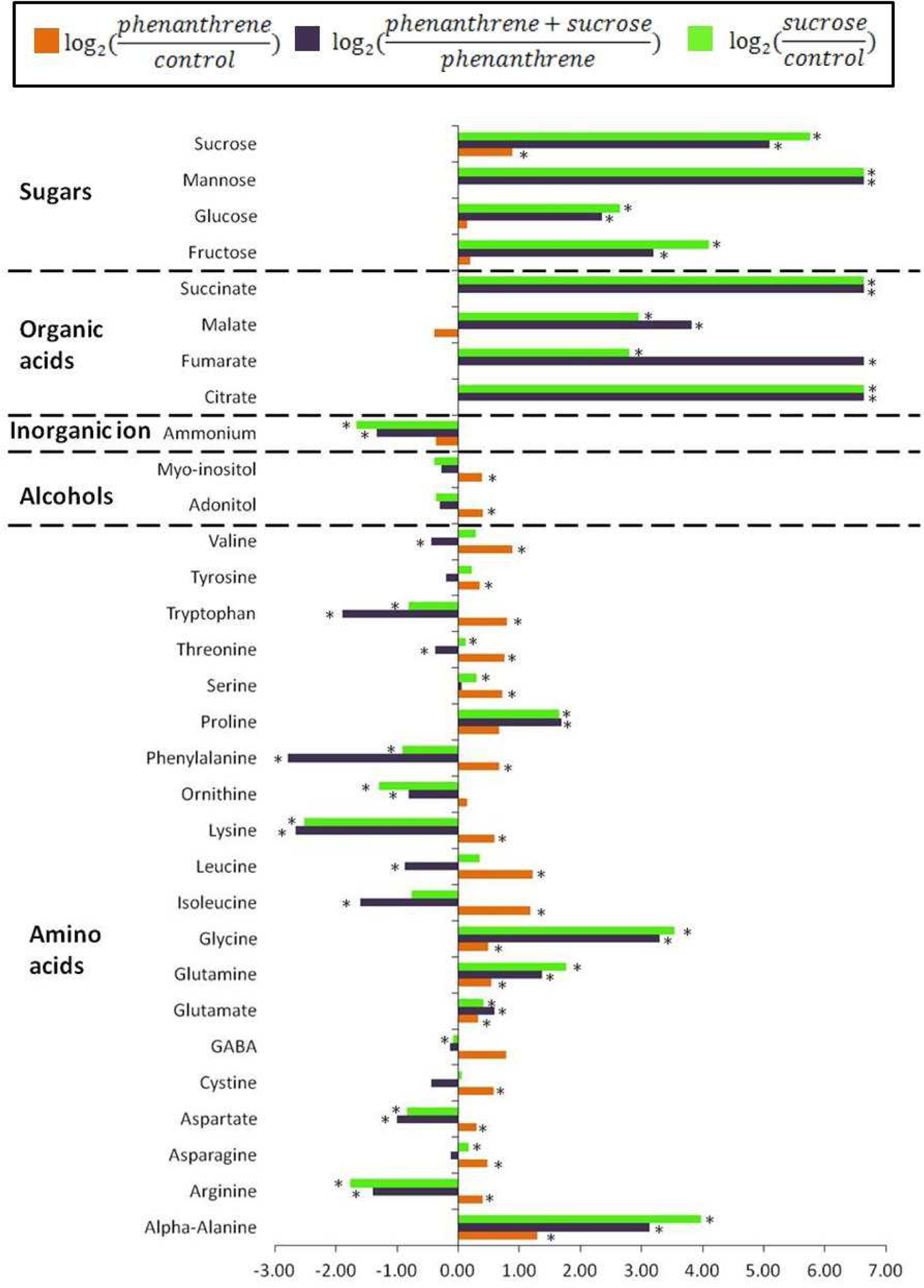
Logarithmic ratios of average metabolites content in whole plants according to the treatment applied. Selected metabolites (with *) are statistically different between both conditions of the ratio (t-Test, p-value≤0.05).

#### 3.4.1. Effect of phenanthrene in unprotected and protected conditions

Figure 11 showed comparison of phenanthrene effects on metabolites ratios of plants grown in different condition. In unprotected condition, 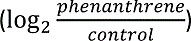, phenanthrene induced significant changes of 20 metabolites levels in comparison to control, while 24 metabolites were impacted by sucrose under phenanthrene induced stress, 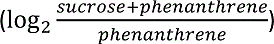

As it was previously observed, phenanthrene induced a slight accumulation of sucrose. However, in protective conditions, we observed an accumulation of not only sucrose but several other sugars (mannose, maltose, glucose, gentiobiose and fructose). Several studies revealed that abiotic stresses such as phenanthrene induced the accumulation of sugars (Gill *et al*., 2003; Dubey and Singh, 1999; Lloyd and Zakhleniuk, 2004). In protective conditions, the accumulation of soluble sugars was highly increased. This result corroborate with the protective effect of the soluble sugars observed in our precedent results and showed in different abiotic stress conditions, such as anoxia (Loreti et al., 2005) and atrazine (Sulmon et al., 2004).

Most of amino acids monitored in our experiment (18 out of 25) increased significantly in non-protective conditions 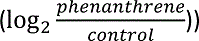. Rai (2002) stated that plants subjected to stress (water, heavy metals, cold, salt…) tended to accumulate amino acids. These authors discussed roles of amino acids that are not only precursors for protein synthesis but also act at different levels (enzyme synthesis and activity regulation, gene expression, redox homeostasis, osmolyte…), this observation may also explain a proteolytic activity related to stressed conditions. Contrary, the comparison of phenanthrene effect in protective against stressed conditions 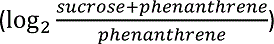 revealed that sucrose induced a deep reconfiguration of metabolites levels as among the 25 amino acids analyzed, 5 amino acids increased (proline, glycine, glutamine, glutamate and alpha-alanine) while 14 amino acids showed significant reduction (Fig. 11).

#### 3.4.2. Sucrose effect on metabolites level is not affected by phenanthrene-induced stress

Figure 11 showed sucrose effects on metabolites ratio in control (green) and stressed condition (purple).

Sucrose induced deep reconfiguration of several metabolites levels in both conditions. 20 metabolites levels were affected by sucrose in non-stressed conditions and 24 in phenanthrene-induced stress conditions. 8 metabolites are affected in only one condition (stressed or control), revealing that phenanthrene did not deeply modify plant response to sucrose.

Sucrose strongly induced the accumulation of sugars showing that plants were able to detect, absorb and metabolize sucrose from their growth media even under phenanthrene induced stress. Sucrose also induced the accumulation of organic acids, which are intermediates of the TCA (see Fig 12). All together, these results revealed a strong regulation of the primary metabolism, suggesting that plants used the sucrose from their growth medium.

**Figure 12:**
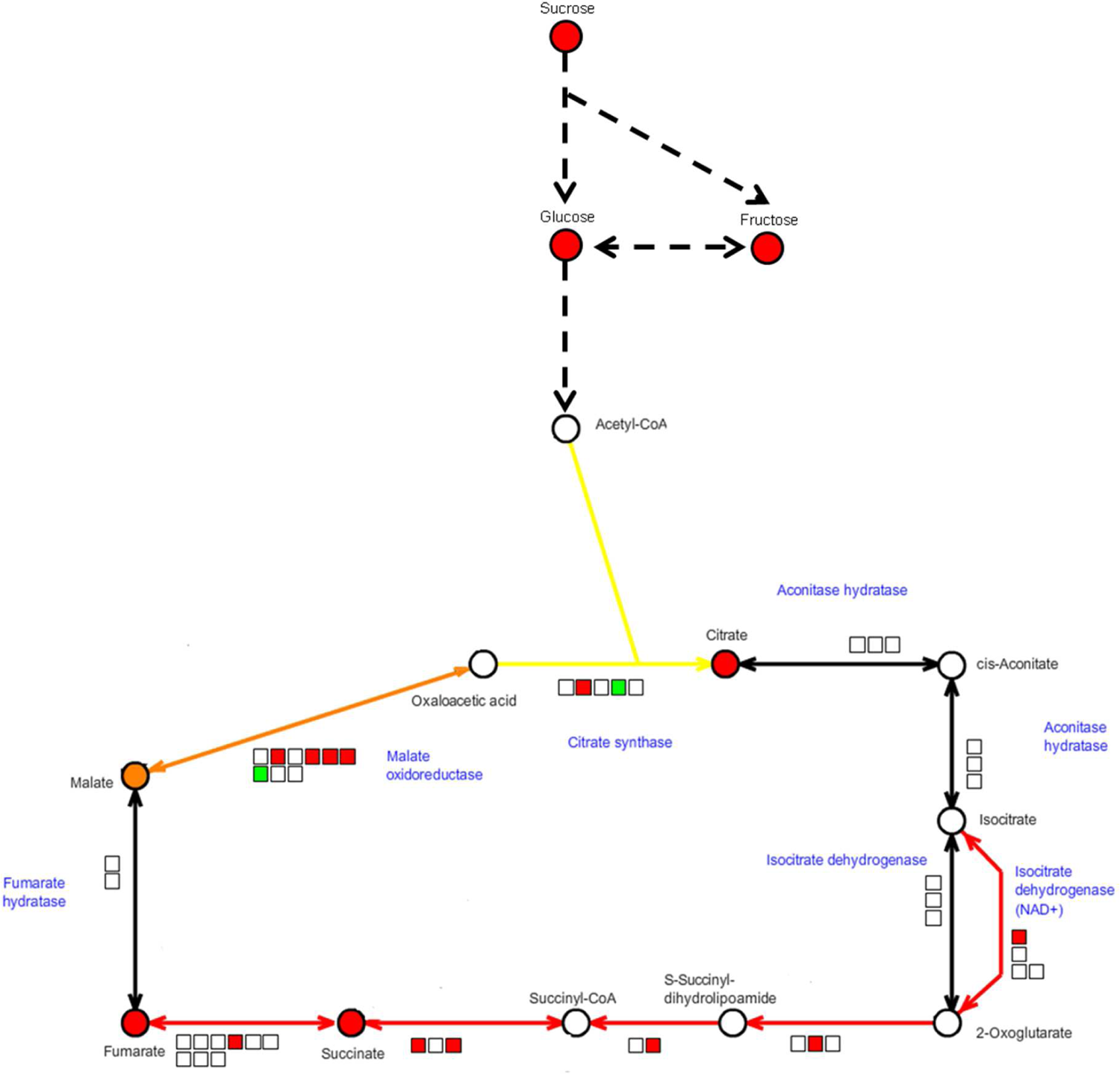
Citric acid cycle and glycolysis representation. Circles showed the metabolomic data from the experiment phenanthrene vs phenanthrene+sucrose (Raw data are in table 1). Squares represented differentially expressed genes from the transcriptomic experiment for an incubation time of 8h. Modified from diagrams acquired with the online software KaPPaView 4 (Tokimatsu et al., 2005)

Metabolomic and transcriptomic changes induced by sucrose under phenanthrene induced-stress revealed a strong activation of primary metabolism suggesting that plants absorb and metabolize through glycolysis and TCA the exogenous sucrose (Fig 12).

In parallel, sucrose decreased ammonium levels in both condition. Amino acids content is also strongly regulated by sucrose, which suggested that nitrogen is stored as amino acids. These changes in nitrogen compounds revealed the impact of the addition of a carbon source on nitrogen metabolism and showed the importance of the balance between nitrogen and carbon (Coruzzi and Zhou, 2001).

## 4. Conclusion

Sucrose enhanced plants development under phenanthrene-induced stress. This effect is specific to sucrose and not due to a change in the osmolarity of the growth medium. Glucose, another sugar with signaling properties, has a limited effect on plant development under phenanthrene-induced stress as it was previously observed for another organic contaminant, the atrazine (Sulmon *et al*., 2004). All these results revealed the specific protective action of sucrose towards phenanthrene. This effect is similar to the one observed with atrazine, but do not confers a complete protection against phenanthrene as it was observed for the herbicide. Transcriptomic and metabolomic data suggested that Arabidopsis absorbed and metabolized sucrose. From these results, the main role of sucrose seemed to be helping the plant to maintain its primary metabolism. Sucrose also showed sensing properties with the strong regulation of carbon and nitrogen metabolism.

Few differences have been observed in changes of metabolites levels induced by sucrose under normal conditions and phenanthrene-induced stress conditions. Phenanthrene induced a lot of changes in metabolites levels of plants grown on medium without sucrose. This effect of phenanthrene is quite limited when plants are grown on medium supplemented with sucrose. Sucrose seemed to counterbalance the effects of phenanthrene on plants metabolism observed in our work on phenanthrene effects (Dumas et al (2016).

Through the activation of genes for enzymes of the redox system, sucrose protected plants from the oxidative damages that might be induced by phenanthrene. Sucrose strongly repressed stress response genes that were induced by phenanthrene, revealing that sucrose mitigated phenanthrene induced stress which allowed the plant to have a better development.

Phenanthrene quantification and cellular localization revealed that plants grown on sucrose and phenanthrene supplemented media accumulated less free-phenanthrene and stored it in different compartments than plants grown on media with phenanthrene only. These results combined by the strong regulation of sucrose on the xenome could reveal a better metabolization of sucrose through the green-liver process.

Adding sucrose in the growth media helped plants to develop in the presence of high doses of phenanthrene. This observation will open new opportunities to improve phytoremediation of PAHs contaminated sites.

---

**Tableau S1:** Complete data from the metabolomic analysis. Numbers in bold are ratios with a <u>statistical (t-Test, p-value≤0.05) difference between both</u> conditions.

|  | Log<sub>2</sub> (phenanthrene/<br>control) | Log<sub>2</sub> (sucrose +<br>phenanthrene/ phenanthrene) | Log<sub>2</sub> (sucrose/<br>control) |
| --- | --- | --- | --- |
| Adonitol | **0.40** | -0.31 | -0.37 |
| Alpha-Alanine | **1.29** | **3.14** | **3.98** |
| Ammonium | -0.37 | **-1.34** | **-1.66** |
| Arginine | **0.40** | **-1.40** | **-1.76** |
| Asparagine | **0.48** | -0.12 | **0.17** |
| Aspartate | **0.29** | **-1.00** | **-0.83** |
| Beta-Alanine | -3.35 | 0.25 | -3.54 |
| Cellobiose | 0.00 | 0.00 | 0.00 |
| Citrate | 0.00 | **6.64** | **6.64** |
| Cystine | **0.58** | -0.44 | 0.06 |
| Fructose | 0.20 | **3.20** | **4.11** |
| Fumarate | 0.00 | **6.64** | **2.80** |
| GABA | 0.79 | -0.14 | **-0.08** |
| Galactinol | 0.00 | 0.00 | 0.00 |
| Galactose | 0.00 | 0.00 | 0.00 |
| Gentiobiose | 0.00 | 0.00 | 0.00 |
| Glucose | 0.14 | **2.35** | **2.65** |
| Glutamate | **0.33** | **0.60** | **0.41** |
| Glutamine | **0.54** | **1.37** | **1.77** |
| Glycerate | 0.00 | 0.00 | 0.00 |
| Glycine | **0.49** | **3.30** | **3.53** |
| Histidine | 0.28 | 0.07 | 0.39 |
| Hydroxyproline | 0.00 | 0.00 | 0.00 |
| Isoleucine | **1.18** | **-1.60** | -0.76 |
| Leucine | **1.22** | **-0.88** | 0.35 |
| Lysine | **0.59** | **-2.66** | **-2.52** |
| Malate | -0.39 | **3.83** | **2.95** |
| Maltose | 0.00 | 0.00 | 0.00 |
| Mannitol | 0.00 | 0.00 | 0.00 |
| Mannose | 0.00 | **6.64** | **6.64** |
| Melibiose | 0.00 | 0.00 | 0.00 |
| Methionine | 0.52 | -0.17 | -0.77 |
| Methylcysteine | -0.85 | -0.39 | -1.13 |
| Myo-inositol | **0.39** | -0.28 | -0.39 |
| Ornithine | 0.15 | **-0.81** | **-1.29** |
| Phenylalanine | **0.67** | **-2.79** | **-0.91** |
| Proline | 0.67 | **1.68** | **1.66** |
| Quinate | 0.00 | 0.00 | 0.00 |
| Serine | **0.72** | 0.05 | **0.30** |
| Sorbitol | 0.00 | 0.00 | 0.00 |
| Succinate | 0.00 | **6.64** | **6.64** |
| Sucrose | **0.88** | **5.09** | **5.76** |
| Threonine | **0.75** | **-0.38** | **0.12** |
| Trehalose | 0.00 | 0.00 | 0.00 |
| Tryptophan | **0.80** | **-1.89** | **-0.81** |
| Tyrosine | **0.35** | -0.19 | 0.22 |
| Valine | **0.89** | **-0.44** | 0.29 |

